# APA-Scan: Detection and Visualization of 3’-UTR Alternative Polyadenylation with RNA-seq and 3’-end-seq Data

**DOI:** 10.1101/2020.02.16.951657

**Authors:** Naima Ahmed Fahmi, Khandakar Tanvir Ahmed, Jae-Woong Chang, Heba Nassereddeen, Deliang Fan, Jeongsik Yong, Wei Zhang

## Abstract

**Background:** The eukaryotic genome is capable of producing multiple isoforms from a gene by alternative polyadenylation (APA) during pre-mRNA processing. APA in the 3’-untranslated region (3’-UTR) of mRNA produces transcripts with shorter or longer 3’-UTR. Often, 3’-UTR serves as a binding platform for microRNAs and RNA-binding proteins, which affect the fate of the mRNA transcript. Thus, 3’-UTR APA is known to modulate translation and provides a mean to regulate gene expression at the post-transcriptional level. Current bioinformatics pipelines have limited capability in profiling 3’-UTR APA events due to incomplete annotations and a low-resolution analyzing power: widely available bioinformatics pipelines do not reference actionable polyadenylation (cleavage) sites but simulate 3’-UTR APA only using RNA-seq read coverage, causing false positive identifications. To overcome these limitations, we developed APA-Scan, a robust program that identifies 3’-UTR APA events and visualizes the RNA-seq short-read coverage with gene annotations.

**Methods:** APA-Scan utilizes either predicted or experimentally validated actionable polyadenylation signals as a reference for polyadenylation sites and calculates the quantity of long and short 3’-UTR transcripts in the RNA-seq data. APA-Scan works in three major steps: (i) calculate the read coverage of the 3’-UTR regions of genes; (ii) identify the potential APA sites and evaluate the significance of the events among two biological conditions; (iii) graphical representation of user specific event with 3’-UTR annotation and read coverage on the 3’-UTR regions. APA-Scan is implemented in Python3. Source code and a comprehensive user’s manual are freely available at https://github.com/compbiolabucf/APA-Scan.

**Result:** APA-Scan was applied to both simulated and real RNA-seq datasets and compared with two widely used baselines DaPars and APAtrap. In simulation APA-Scan significantly improved the accuracy of 3’-UTR APA identification compared to the other baselines. The performance of APA-Scan was also validated by 3’-end-seq data and qPCR on mouse embryonic fibroblast cells. The experiments confirm that APA-Scan can detect unannotated 3’ -UTR APA events and improve genome annotation.

**Conclusion:** APA-Scan is a comprehensive computational pipeline to detect transcriptome-wide 3’-UTR APA events. The pipeline integrates both RNA-seq and 3’-end-seq data information and can efficiently identify the significant events with a high-resolution short reads coverage plots.

## Introduction

Poly(A)-tails are added to pre-mRNA after the polyadenylation signal (PAS) during the 3’-end processing of pre-mRNA [1]. The last exon of mRNA contains a non-coding region, 3’-untranslated region (3’-UTR), which spans from the termination codon to the polyadenylation site. The 3’-UTR acts as a molecular scaffold to bind microRNAs and RNA-binding proteins and functions in regulatory gene expression [2]. In human and mouse, more than 70% of genes contain multiple PASs in their 3’ - UTRs and polyadenylation using upstream PASs leads to the production of mRNA with shortened 3’-UTRs (3’-UTR APA) [3, 4]. 3’-UTR APA is known to increase the efficiency of translation and is associated with T cell activation, oncogene activation, and poor prognosis in many diseases [5, 6, 7]. Recent study has demonstrated that 3’ - UTR APA is one way to increase protein synthesis without increasing the quantities of mRNAs, indicating that it is an important element in gene expression which cannot be understood by conventional differential gene or transcript expression analysis [8]. Up-regulation of mTOR signaling pathway can lead to transcriptome-wide 3’-UTR APA [8, 9].

3’-UTR APA has gained much attention recently and the importance of the 3’ - UTR APA in human diseases has been demonstrated as mentioned above. Some recent studies show that both proliferating cells and transformed cells favor expression of shorter 3’-UTR through APA and lead to the activation of oncogenes [6, 10]. Some other research shows the trend in cancer cells for highly expressed genes to exhibit shorter 3’-UTR with fewer microRNA binding sites, decreasing microRNA-mediated translation repression [5, 11]. All these studies imply that 3’-UTR APA may serve as a new layer of prognostic biomarker. A scalable computational model is highly needed to detect the genome-wide unannotated 3’-UTR APA in different phenotypes.

Several bioinformatics pipelines are available for the analysis of UTR-APA using RNA-seq data [12, 13, 14, 15, 16]. In general, all these methods measure the changes in 3’-UTR lengths by modeling the RNA-seq read density change near the 3’-end of mRNAs. Indeed, with the aid of these methods, RNA-seq experiments became a powerful approach to investigate 3’-UTR APA. However, in many cases the identified APA sites are not functionally and physiologically relevant because most pipelines do not reference actionable PASs in their 3’-UTR APA simulation. RNA-seq is not particularly accurate when it comes to identifying polyadenylation sites, making novel APA transcript identification rather difficult. Therefore, 3’-end-seq data has been developed to address these issues by enriching for 3’-end reads in high-throughput sequencing experiment [17] and provides the accurate polyadenylation sites. In addition to the limitations of the current bioinformatics pipelines mentioned above, none of them can provide high-resolution read coverage plots of the APA events with an accurate annotation. We have developed APA-Scan (Figure 1), a bioinformatics program, to detect and visualize genome-wide 3’-UTR APA events. APA-Scan integrates both 3’-end-seq data and the location information of predicted canonical PASs with RNA-seq data to improve the quantitative definition of genome-wide UTR APA events. APA-Scan efficiently manages large-scale alignment files and generates a comprehensive analysis for UTR APA events. It is also advantageous in producing high quality plots of the events.

**Figure 1.**
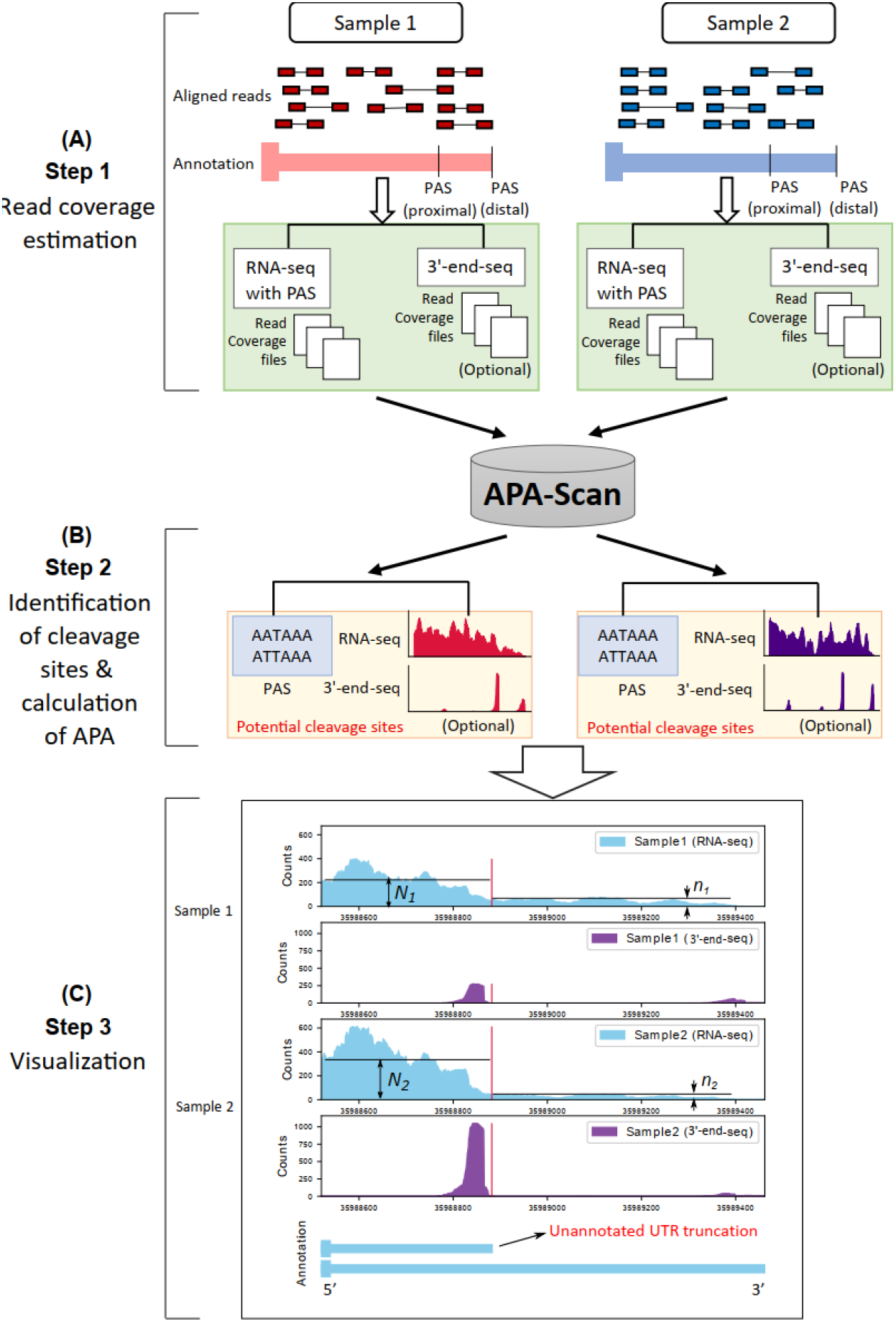
Workflow of APA-Scan. Starting with aligned RNA-seq and 3’-end-seq (optional) bam files, APA-Scan consists of three steps and generates high quality graphical illustration of aligned sequences with the indication of 3’-UTR APA events. (A) Read coverage files are generated for RNA-seq and 3’-end-seq (if provided) input samples. (B) APA-Scan identifies potential cleavage sites according to polyadenylation signal (PAS) hexamer: ATTAAA or AATAAA, or 3’-end peaks (if 3’-end-seq data is available). (C) Graphical illustration of the identified events. The illustration also highlights unannotated short 3’-UTR transcript identified from this task. The vertical red lines show the corresponding cleavage sites.

## Results

APA-Scan is designed to identify both annotated and de novo 3 ‘ -UTR APA events between different biological conditions. To access the performance of APA-Scan, it was compared with two baseline methods on both simulated and real RNA-seq datasets. In the simulation experiment, we first generated synthetic dataset with pre-defined 3’-UTR APA events (ground truth) to test if the APA-Scan and baseline methods can detect them. Next, we performed experiments on two mouse embryonic fibroblast (MEF) cells to evaluate the performance of APA-Scan. The results of analyzing real MEF RNA-seq datasets were validated using both qPCR and 3’-end-seq data.

### Experimental results with simulated RNA-seq data

In the simulation experiment, we generated synthetic RNA-seq short reads with flux-simulator [18]. 1000 pre-defined 3’-UTR APA events were simulated as the ground truth between two different conditions. In each condition, three technical replicates were generated by repeating the experiment three times with the same parameter setting in the flux simulator. The details of the parameters used in this experiment are provided in the Additional file 2. For both conditions, the gene expressions were sampled from a Poisson distribution to reflect a real RNA-seq data [19]. For each gene, one proximal polyadenylation site was synthesized to represent the end of the short isoform and the end of the annotated transcript was applied to define the end of the long isoform of that gene. To generate the ground truth profile of the 3’-UTR APA events, the expression proportions of the short and long isoforms in the same gene were assigned significantly different values in two conditions (i.e., the proportion difference was larger than 10%) to represent the existence of the APA event.

In the simulation experiment, two sets of synthetic data were generated by flux-simulator. One with 30M (30 million) paired-end reads in each replicate and one with 50M paired-end read. In both cases, the read length is 76 bps of each end. APA-Scan was compared with DaPars and APAtrap on the simulated RNA-seq datasets. To detect the significant 3’-UTR APA events, APA-Scan used p-value < 0.05 (χ2-test) as the cutoff. DaPars identified APA events according to the difference in PDUI (Percentage of Distal polyA Usage Index) values between two conditions > 0.1 and FDR < 0.05; whereas APAtrap selected events using the cutoff values of two parameters: percentage difference of APA site usage between two conditions > 0.1 and FDR < 0.05. The performance of the methods is then evaluated using AUC score, sensitivity and specificity. Figure 2 shows that, APA-Scan outperformed the two baselines in terms of AUC scores and got the best score of 0.94 in both sequence depths (30M reads and 50M reads) and followed by APAtrap (0.73 in 30 million reads case and 0.75 in 50 million reads case). DaPars did not work very well compared to the other two methods and the AUC scores were below 0.7 in both cases, though there was an improvement in the case with more reads. We also report the sensitivity and specificity for each method with two different sequencing depths in Table 2. APA-Scan gets the highest sensitivity and specificity scores for both cases, which indicates that APA-Scan outperformed the baseline methods in detecting the true 3’-UTR APA events and eliminating the true negative ones.

**Figure 2.**
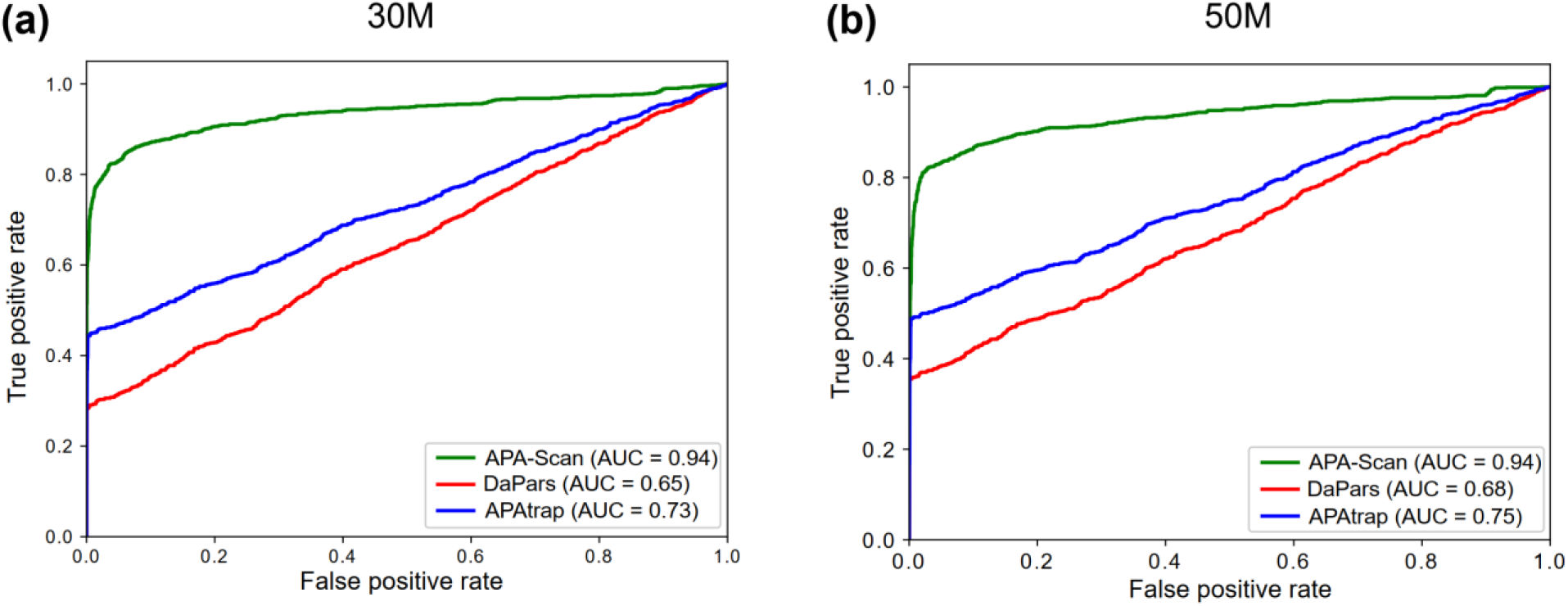
Simulation experiment to assess the performance of APA-Scan and the baseline methods (DaPars and APAtrap). (a) Results on the simulation experiment with 30 million (30M) short reads. (b) Results on the simulation experiment with 50M short reads. The receiver operating characteristic (ROC) curves, i.e., true positive rate against false positive rate, are plotted.

**Table 1.** Categorized overview of the technical parameters of APA-Scan

| Categories | Types | Description |
| --- | --- | --- |
| Input data | APA-Scan<sup>peaks</sup> | Both 3'-end-seq and RNA-seq data are provided |
|  | APA-Scan<sup>PAS</sup> | Only RNA-seq data is provided |
| Mode | Default | Search for shorter 3'-UTR APA events |
|  | Extended | Search for longer 3'-UTR APA events |
| Reported list | Default | List the most significant cleavage site for each gene |
|  | All | List all candidate cleavage sites for each gene |

**Table 2.** Comparison among APA-Scan, DaPars and APAtrap on simulated RNA-seq data with two different sequencing depths (30 million reads and 50 million reads). AUC (the area under the ROC curve) score, sensitivity, specificity of the three methods are reported. The best results across the three methods are bold.

| Reads | Method | AUC | Sensitivity | Specificity |
| --- | --- | --- | --- | --- |
|  | APA-Scan | **0.94** | **0.83** | **0.95** |
| 30M | DaPars | 0.65 | 0.33 | 0.58 |
|  | APAtrap | 0.73 | 0.57 | 0.90 |
|  | APA-Scan | **0.94** | **0.83** | **0.95** |
| 50M | DaPars | 0.68 | 0.41 | 0.71 |
|  | APAtrap | 0.75 | 0.63 | 0.91 |

As different sequencing depths may affect the performance of APA-Scan, we generated five simulation experiments with different read depths, i.e., 2M, 5M, 10M, 30M, and 50M paired-end reads by flux-simulator with the same parameter setting to learn the impact of sequencing depths in the analysis of 3’-UTR APA with APA-Scan. In this experiment, the read length was also 76 bps for each end and three replicates were generated for each condition in each read depth using the same procedures as mentioned in the previous section. Figure 3 shows the ROC curves for different sequencing depth on detecting the 3’-UTR APA events. APA-Scan shows moderate performance with low sequencing depths (i.e., 2M and 5M). However, the performance of APA-Scan improved drastically (AUC = 0.94) after it reached to a certain sequencing depth (i.e., 10M in this study) and holds that performance across read depths above that threshold. This result suggests that APA-Scan is quite robust in detecting APA events on lowly expressed genes and relatively low read coverage samples.

**Figure 3.**
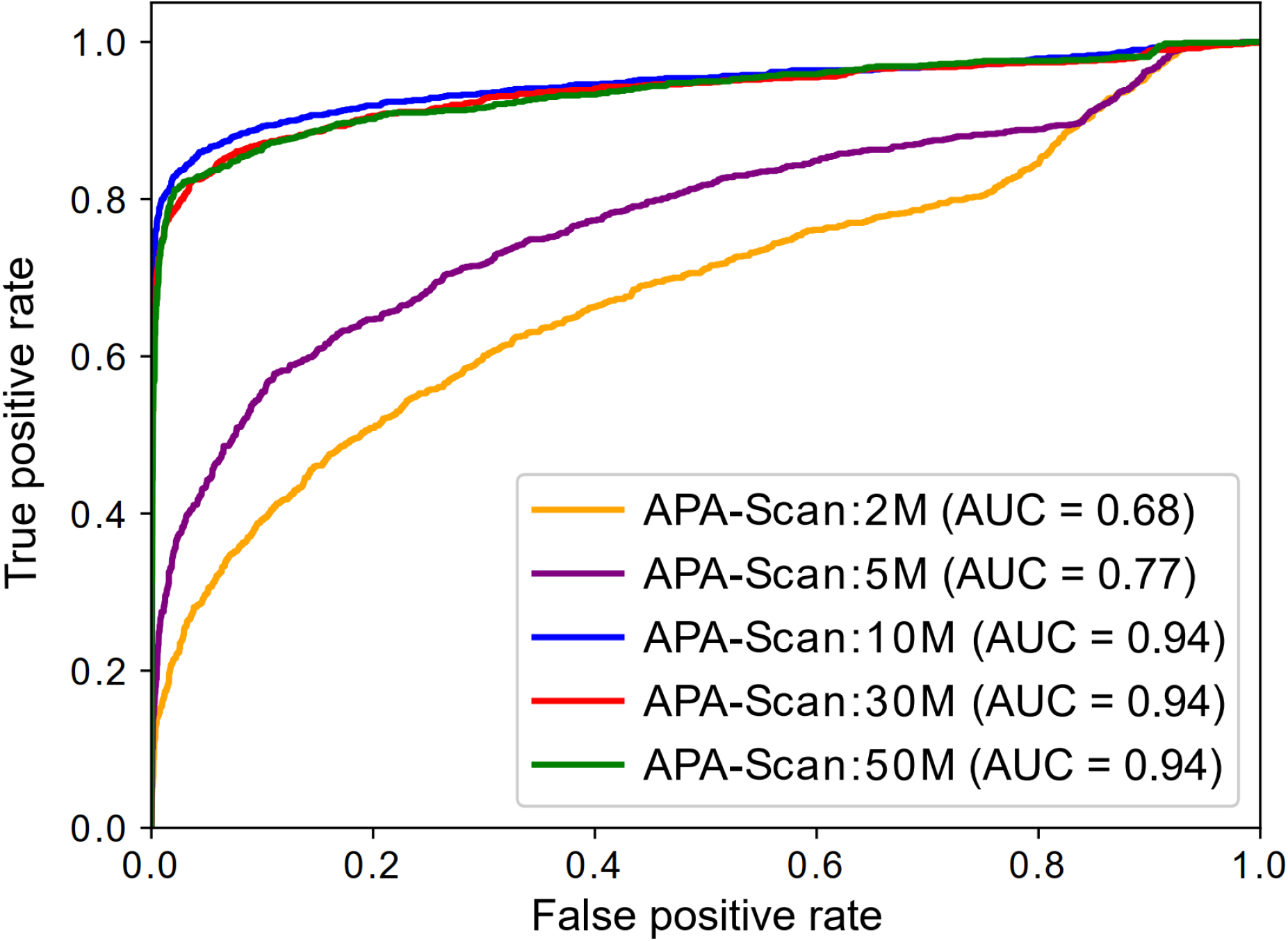
Simulation experiment to assess the performance of APA-Scan on different sequencing depths. The ROC curves for the results of different RNA-seq read depth are plotted.

### Experimental results with MEFs samples

In the real RNA-seq experiments, two MEFs samples Tsc1^-/-^ and WT were used in the analysis to evaluate the performance of APA-Scan and baseline methods. Knockout of Tsc1, a negative regulator of mTOR pathway, leads to uncontrolled mTOR hyper-activation compared with WT. For the comparison and evaluation purposes, the APA-Scan was run on two different setups. One used PASs in the 3’-UTRs as potential cleavage sites, and we denote it by APA-Scan^*PAS*^. The other one considered 3’-end-seq peaks as candidate sites, and it is denoted as APA-*Scan^peaks^.* First, APA-Scan^*PAS*^ was applied to detect 3’-UTR APA events between the two MEFs samples with p-value < 0.05. APA-Scan^*PAS*^ detected 265 events, whereas DaPars and APAtrap detected 785 and 1130 significant events, respectively. These events were then verified by the polyadenylation sites reported by 3’-end-seq data. If a predicted 3’-UTR APA event is within 50 bps upstream or downstream of the loci of the peak(s) in 3’-end-seq data, then this APA event is considered overlapping with the 3’-end-seq signals. Though APA-Scan^*PAS*^ detected less number of significant events compared to the baseline methods, 87.92% (233) of the events were validated by the 3’-end-seq signals according to the result shown in Figure 4 and Table 3. DaPars and APAtrap identified more events than APA-Scan^*PAS*^, however, both the number and ratio of the overlapping events with the 3’-end-seq signals are significantly lower than the events detected by APA-Scan^*PAS*^. Note that APA-Scan^*PAS*^ did not use any information from 3’-end-seq data to identify the APA events. These results concur with our findings in the simulation experiment that APA-Scan not only do better detection on the true APA events but also prevent the false positives. Figure 5 shows the number of overlapped genes with the 3’-UTR APA events detected by the three methods. From the results, we can conclude that the agreement of the three methods is not high and most identified events were only detected by one method.

**Figure 4.**
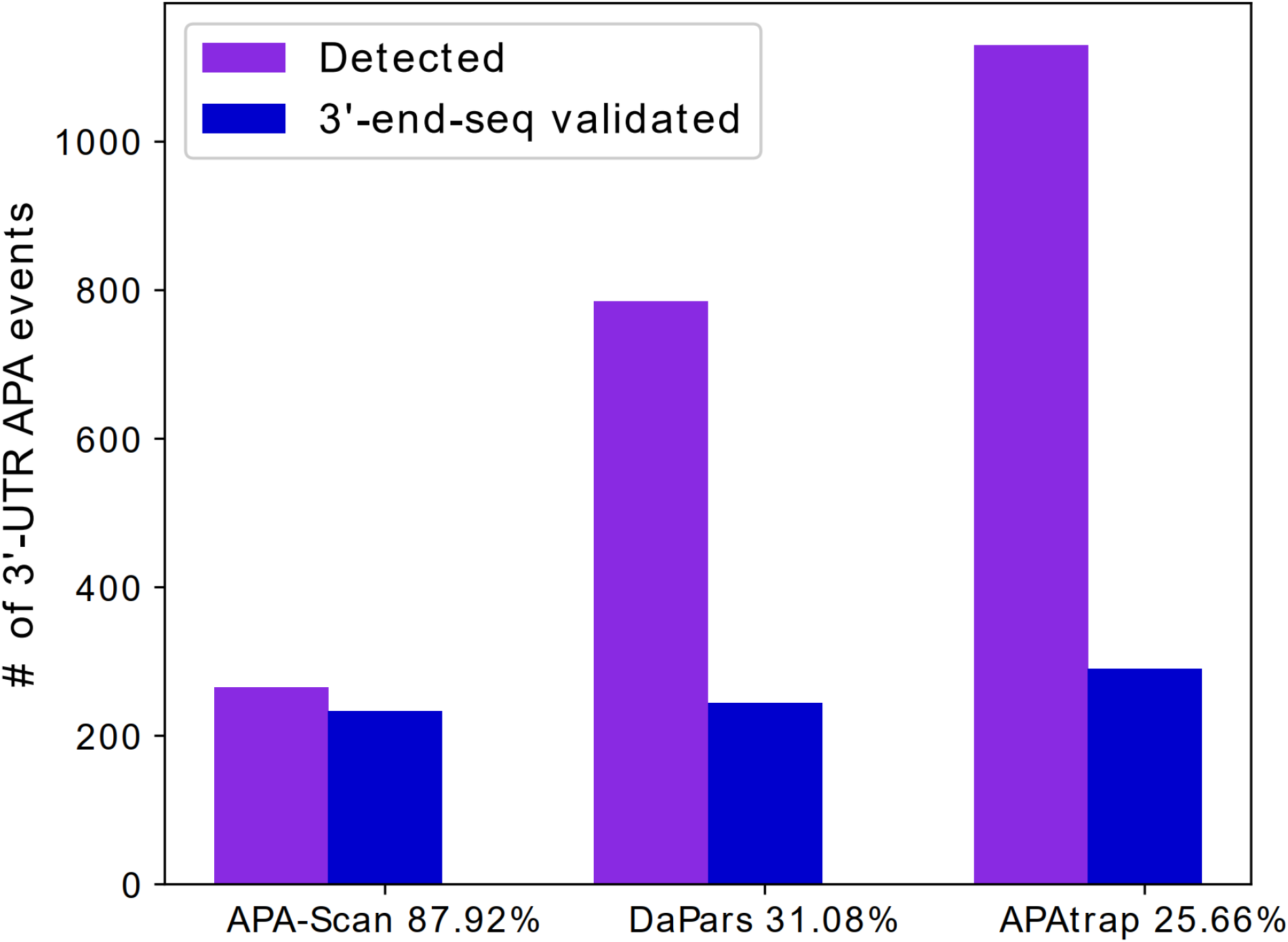
Evidence of polyadenylation sites supported by 3’-end-seq data for 3’-UTR APA events detected by different methods in MEFs samples. The number of events predicted by each method are shown in purple and the number of events validated by the signals in the 3’-end-seq data are shown in blue. The x-axis shows the percentage of the identified events is validated by 3’-end-seq.

**Figure 5.**
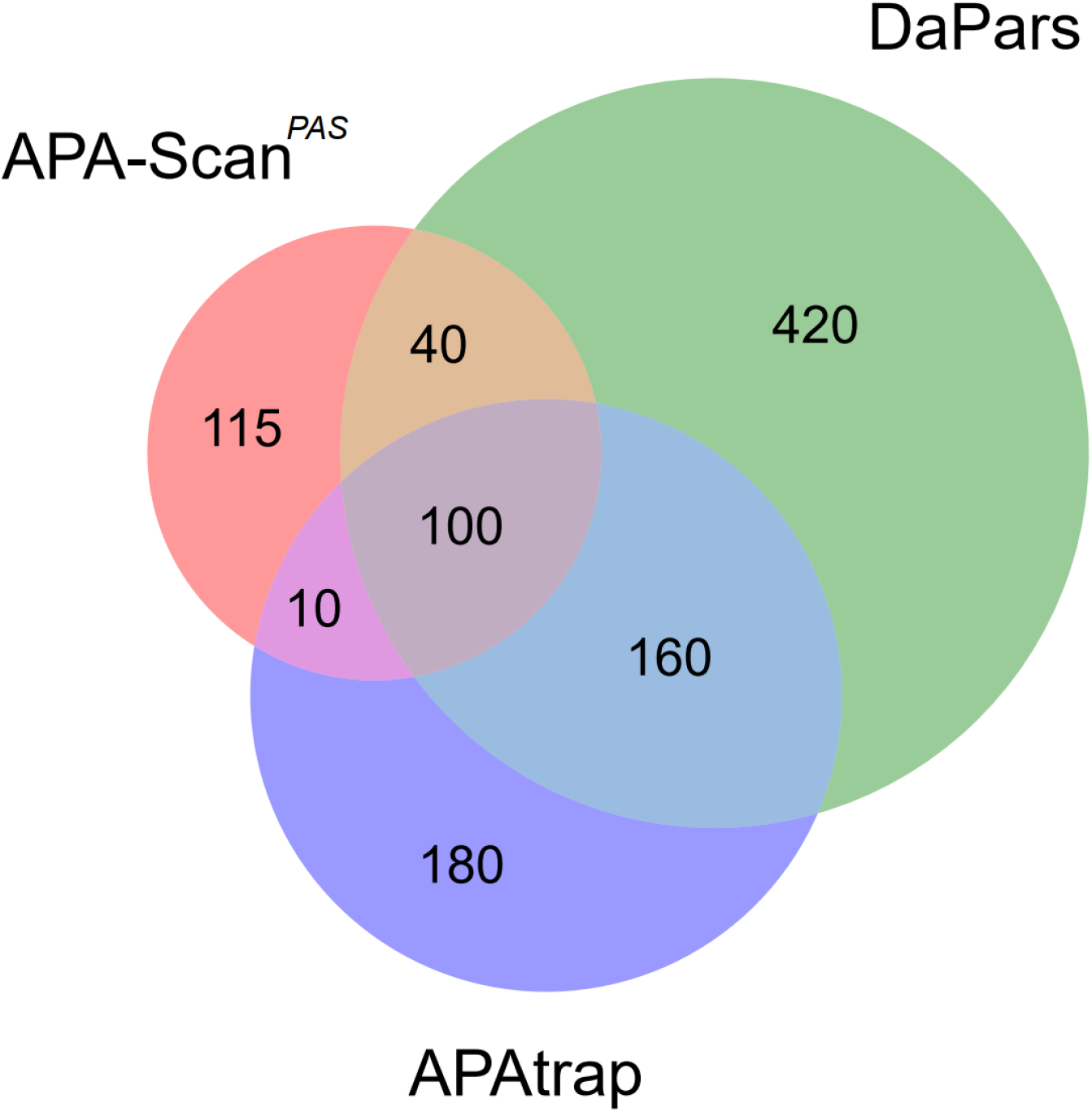
Venn diagram shows the overlapped genes with the 3’-UTR APA events identified by three methods (i.e., APA-Scan, DaPars and APAtrap) between two MEFs samples (WT vs Tsc1^-/-^).

**Table 3.** Number of events detected by APA-Scan, DaPars and APAtrap and validated by 3’-end-seq data for MEFs samples.

| Method | Detected | Validated by 3'-end-seq | Ratio (%) |
| --- | --- | --- | --- |
| APA-Scan | 265 | 233 | 87.92 |
| DaPars | 785 | 244 | 31.08 |
| APAtrap | 1130 | 290 | 25.66 |

To further validate the analysis results by APA-Scan, we conducted qPCR experiments for Srsf3 and Rpl22 transcripts from Tsc1^-/-^ and WT MEFs based on the significant 3’-UTR APA events reported by APA-Scan^*peaks*^. These genes were selected due to the design of PCR (polymerase chain reaction) primers for wet-lab validation. As shown in Figure 6, both Srsf3 and Rpl22 showed the increase of the short 3’-UTR transcript by APA in Tsc1^-/-^ compared to WT MEFs, which is consistent with our observations on the RNA-seq and 3’-end-seq read coverage plots. These results further confirm that APA-Scan can identify the true 3’-UTR APA events with RNA-seq and 3’-end-seq samples from two different biological contexts. The more details of the qPCR analysis and the primer sequences of the two genes are available in the Additional file 1.

**Figure 6.**
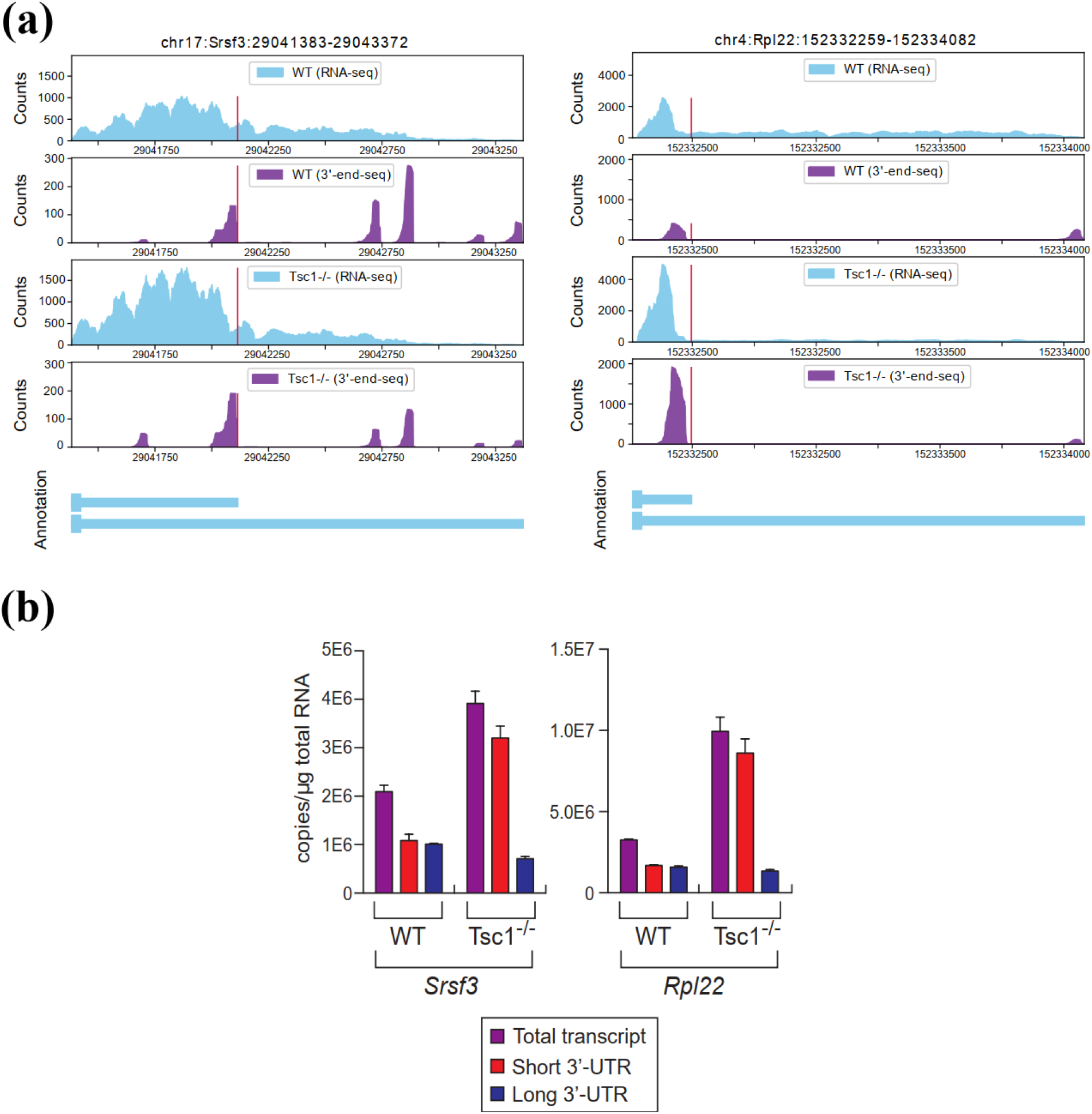
Experimental results: (a) RNA-seq and 3’-end-seq read coverage plots of the 3’-UTR in *Srsf3* and *Rpl22* gene in the two samples with isoform annotation. (b) The level of total, short 3’-UTR, and long 3’-UTR transcripts from *Srsf3* and *Rpl22* was measured by qPCR. Because it is not possible to design specific primers for the qPCR analysis of short 3’-UTR transcript, the amount of short 3’-UTR transcripts were calculated by subtracting the quantity of long 3’-UTR transcripts from total.

Generally, the nucleotide profiles surrounding the polyadenylation sites are dominated by two motifs and their variants: AATAAA and ATTAAA and these two hexamers are observed upstream of the cleavage sites [20]. This phenomenon leads us to explore the nucleotide composition near the predicted polyadenylation sites by APA-Scan^*peaks*^. Figure 7 shows a high concentration/cluster of nucleotide ‘A’ in the polyadenylation site, positioned at 0. The upstream surrounding region is also dominated by ‘A’ and ‘T’, which clearly indicates the existence of potential 3’-UTR APA events.

**Figure 7.**
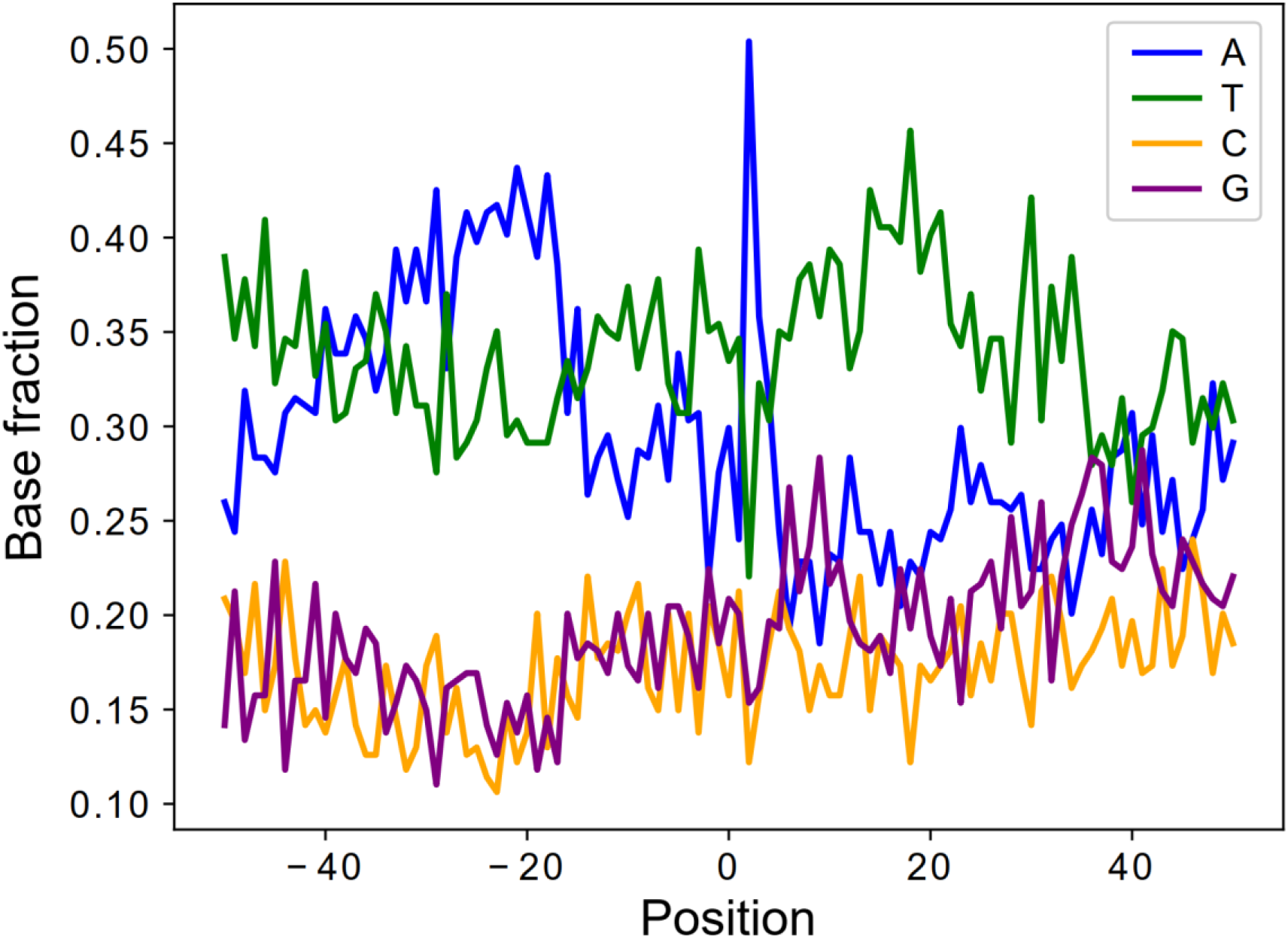
Nucleotide composition of the sequence surrounding the polyadenylation sites identified by APA-Scan^*peaks*^ for MEFs samples. 50bp up and downstream region is plotted with base sequences. x-axis denotes the position in the region, 0 is the location of the identified polyadenylation site. y-axis shows the fraction of the nucleotides content at each position.

## Discussion

APA is one mechanism for post-transcriptional regulation of mRNA expression, and it is defined as use of more than one polyadenylation sites. 3’-UTR APA is one of the most frequent APA forms, which contains more than one polyadenylation sites in the 3’-UTR. It generates multiple mRNA transcripts with different 3’-UTR lengths without affecting the protein encoded by the gene. Since the 3’-UTR of mRNA of-ten contains binding sites for microRNAs, 3’-UTR APA potentially leads to altered mRNA stability or protein translation efficiency due to variation of 3’-UTR length. Identification and assessment of APA sites has been a major goal in understanding transcriptomic diversity. Several bioinformatics tools have been developed to predict transcriptome-wide polyadenylation sites with RNA-seq data. However, our experimental results on simulated and real samples indicate that the current methods (e.g., DaPars and APAtrap) can detect large number of APA events, but significant portion of the events are false positives. A similar data analysis on BT549 breast cancer cells (mock vs. torin 1 treated) in Figure S1 in the Additional file 1 illustrates a similar pattern. By integrating 3’-end-seq and RNA-seq data, APA-Scan can potentially reduce the number of false positive events. To evaluate the performance of APA-Scan on real cancer patient samples, one pair (tumor vs. matched normal tissue) of The Cancer Genome Atlas (TCGA) breast cancer samples are also analyzed and reported in the Additional file 1. Figure S2 shows that a significant portion (>72%) of the 3’-UTR APA events are not differentially expressed. There-fore, APA-based molecular signatures could provide additional predictive power of cancer outcomes by combining the differently expressed genes.

APA-Scan not only can accurately detect the splicing events compared to the baseline methods, but also provides reasonable running time. Table 4 shows a com- parison of the CPU time of each method on Tsc1^-/-^ and WT MEFs. The CPU time was measured on an Intel(R) Xeon(R) CPU E5-2620 v4 @ 2.10GHz machine. Both APA-Scan and APAtrap completed the analysis in a similar amount of time. However, DaPars is much slower than the other two methods which is not suit- able to be applied on large-scale experiment in terms of running time. Overall, this study reports an efficient and precise framework for 3’-UTR APA identification with RNA-seq and 3’-end-seq data.

**Table 4.** CPU time of APA-Scan, DaPars and APAtrap on two MEFs samples (WT vs Tsc1^-/-^).

| APA-Scan | DaPars | APAtrap |
| --- | --- | --- |
| 48 mins | 21 hours | 36 mins |

## Conclusion

We developed APA-Scan, which offers a comprehensive computational pipeline to identify transcriptome-wide 3’-UTR APA events. By integrating RNA-seq data and 3’-end-seq information (experimentally verified or computationally predicted). APA-Scan can efficiently identify significant APA events and also, can illustrate the events with read coverage plots. 3’-end-seq signals and the wet-lab experiment using qPCR demonstrate that APA-Scan provides high-accuracy and quantitative profiling of 3’-UTR APA events. Therefore, we expect that, APA-Scan will serve as a useful tool for APA site analysis.

## Methods

### APA-Scan pipeline

APA-Scan workflow comprises of three major steps: (i) read coverage estimation; (ii) identification of polyadenylation sites and the calculation of APA; (iii) graphical illustration of UTR APA events (Figure 1). First, APA-Scan takes aligned RNA- seq and 3’-end-seq data from two different biological conditions as input. Each biological condition can have multiples samples or replicates. The read coverage files are generated by SAMtools [21]. In this step, the 3’-end-seq data is an optional input.

In the second step, APA-Scan starts the analysis by extracting 3’-UTR frames for each gene. APA-Scan is designed in two modes: (a) Default, and (b) Extended. All the aligned reads from 3’-end-seq data are pooled together to identify peaks and the corresponding unannotated cleavage sites in 3’-UTR regions and downstream of the 3’-UTR regions (i.e., APA-Scan^*peaks*^). In the Default mode, 3’-UTR regions are selected according to the end of the longest annotated transcript of the gene. The loci of peaks identified in the 3’-end-seq data are considered as potential cleavage sites. If the 3’-end-seq data is not provided by the user, detected PASs (generally two variations of the hexamers: AATAAA, ATTAAA) in 3’-UTRs are considered as the potential cleavage sites (i.e., APA-Scan^*PAS*^) follow the ideas in Omni-PolyA [16] which use 12 most common PAS variants to determine the cleavage sites. In the Extended mode of APA-Scan, the potential peaks/PAS signals are searched up to 10kb downstream of the end of transcript to discover de novo distal polyadenylation sites. The locations detected from all input samples are merged to get a combined list of potential cleavage sites. The major commands and general terminologies to run APA-Scan are listed in Table 1.

APA-Scan evaluates each empirical cleavage site in the 3’-UTR of a transcript by contrasting the RNA-seq short reads coverage up and downstream of the candidate site between the two biological conditions. n and N denote the average read coverage up and downstream of the site. They are determined by estimating the number of reads mapped to upstream and downstream of the cleavage site, *r_u_* and *r_d_*, divided by their effective length, *l_u_* and *l_d_*, respectively (i.e., 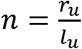 and 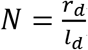). For each potential polyadenylation site, the ratio differences between the samples in two conditions are calculated based on the following equation

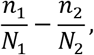

where 1 and 2 represent the two conditions. Ratio difference indicates the change in read coverage between two conditions and only the absolute ratio difference > 0.1 is considered as candidate site for the further analysis. After that, the canonical 2 x 2 χ2-test is applied to report the p-value for each candidate site. The χ2-test measures how much the observation deviates from the null hypothesis. In our experiment, we set the null hypothesis as the average read coverage before and after the cleavage sites are consistent among the two biological conditions. For any true 3’-UTR APA event, there must be a significant read coverage drop-off around the cleavage sites, and the ratios of the average read coverages before and after the cleavage sites are crucially different in the two conditions. In such cases, the χ2-test precisely reports significant p-values to reject our null hypothesis. APA-Scan will report both significant and insignificant in an Excel file. A comprehensive user’s manual is provided in the Additional file 2.

In the third step, based on the significance of 3’ -UTR APA events calculated in the previous step, APA-Scan generates RNA-seq and 3’-end-seq (if provided) read coverage plots with the 3’-UTR annotations for one or more user-specific events. Users may specify the region of the genome locus to generate the read alignment plot. Figure 1 (Step 3) illustrates an example of the read coverage plot generated by APA-Scan.

### Baselines and evaluation methods

In this study, two widely used 3’-UTR APA identification approaches, DaPars [12] and APAtrap [15] were applied to compare the performance with APA-Scan. The command lines to run the baseline methods are available in the Additional file 1. To evaluate the performance of APA-Scan and baseline methods, the area under the ROC curve (AUC), sensitivity and specificity were used on the identified lists of 3’-UTR APA events.

### Short read alignments and peak identification

In this study, two mouse embryonic fibroblasts (MEFs) samples and two breast cancer cell lines (BT549) were used in the analysis to evaluate the performance of APA-Scan and baseline methods. For the MEFs samples, we performed RNA-seq and 3’-end-seq analyses of poly(A+) RNAs isolated from Tsc1^-/-^ and wild-type (WT) MEFs. In the RNA-seq analysis, 63,742,790 paired-end reads for WT and 74,251,891 paired-end reads for Tsc1^-/-^ MEFs were produced from Hi-Seq pipeline with length of 50 bps of each end. The short reads were aligned to the mm10 reference genome by TopHat2 [22], allowing up to two mismatches. Finally, 87.1% of short reads from WT and 87.5% of sequence reads from Tsc1^-/-^ MEFs were mapped to the reference genome for APA analysis in the study. In the 3’-end-seq analysis, the reads from WT and Tsc1^-/-^ MEFs were preprocessed to trim A’s off the 3’-ends and then filtered by removing the reads of low-quality 3’-end (Phred score < 30) and shorter than 25 bps. The remaining reads were aligned to the mm10 reference genome by Bowtie [23] without allowing any mismatches. In total, 6,186,893 paired-end reads were aligned for WT and 5,382,111 reads were aligned for Tsc1^-/-^. All aligned reads from 3’-end-seq were pooled together in order to identify peaks and the corresponding cleavage sites in the reference genome by the read coverage signals. In each read alignment ‘hill’, the location with the highest read coverage between two zero coverage positions was considered as the peak of the ‘hill’. The 3’-end of the peak is chosen as the potential corresponding cleavage sites where the read coverage at the peak quantifies the cleavage at the site. For the breast cancer cell lines, we performed RNA-seq analysis of poly(A+) RNAs isolated from BT549 mock and Torin1 treated cells. 131,955,082 paired-end reads for BT549 mock, and 138,127,113 paired-end reads for BT549 treated with Torin1 were produced from Hi-Seq pipeline with length of 51 bps of each end. The short reads were aligned to the hg38 reference genome by TopHat2, allowing up to two mismatches. Finally, 85.2% of short reads from BT549 mock and 84.7% of sequence reads from BT549 treated with Torin1 were mapped to the reference genome for APA analysis in the study.

## Declarations

## Abbreviations

APA: alternative polyadenylation
3’-UTR: 3’-untranslated region
mTOR: mechanistic target of rapamycin
PAS: polyadenylation signal
MEFs: mouse embryonic fibroblasts
WT: wild-type
ROC: receiver operating characteristic
AUC: area under the ROC curve
qPCR: quantitative polymerase chain reaction

## Ethics approval and consent to participate

Not applicable.

## Consent for publication

Not applicable.

## Availability of data and materials

The source code in this study is available at: https://github.com/compbiolabucf/APA-Scan. The accession number for the MEFs RNA-seq data in this study is <u>SRP056624</u>. The accession number for the 3’-end-seq data in this study is <u>SRP133833</u>.

## Author’s contributions

NAF, DF, JY, and WZ conceived the study and planned the analysis. NAF, KTA, and HN performed data analysis. JWC and JY designed and performed qPCR experiments. NAF, KTA, JY, and WZ wrote the manuscript. All authors read and approved the final manuscript.

## Additional Files

### Additional file 1

Figure S1; Figure S2; The command lines used for running the baseline methods; Parameters to run flux-simulator; qPCR analysis and primer sequences.

### Additional file 2

User’s manual of APA-Scan

## Supplementary

### 1 Running the baselines

#### 1.1 DaPars

Inputs:

- BED files. Flux-simulator simulated three fastq files for three replicates in each condition. Using SAMtools (v0.1.8), read coverage files for each chromosome are generated in BAM format from the fastq files. Six bedgraph files in two conditions are generated from the BAM files using BEDtools.
- Gene annotation in .bed format: mm10_Refseq.bed
- Configuration file: configure.txt Annotated_3UTR=mm10_Refseq_extracted_3UTR.bed Group1_Tophat_aligned_Wig = case_1.bedgraph, case_2.bedgraph, case_3.bedgraph Group2_Tophat_aligned_Wig = control_1.bedgraph, control_2.bedgraph, control_3.bedgraph Output_directory = DaPars_out/ Output_result_file = Dapars_out Num_least_in_group1 = 1 Num_least_in_group2 = 1 Coverage_cutoff = 30 FDR_cutoff = 0.05 PDUI_cutoff = 0.1 Fold_change_cutoff = 0.59

**Command 1:** python DaPars_Extract_Anno.py -b mm10_Refseq.bed -s mm10_Refseq_id.txt -o mm10_Refseq_extracted_3UTR.bed
**Command 2:** python DaPars_main.py configure.txt

#### 1.2 APAtrap

Inputs:

- BED files. Flux-simulator simulated three fastq files for three replicates in each condition. Using SAMtools (v0.1.8), read coverage files for each chromosome are generated in BAM format from the fastq files. Six bedgraph files in two conditions are generated from the BAM files using BEDtools.
- Gene annotation in .bed format: mm10_Refseq.bed

**Command 1:** identifyDistal3UTR -i A1.bedgraph A2.bedgraph A3.bedgraph B1.bedgraph B2.bedgraph B3.bedgraph -m mm10_Refseq.bed -o mm10.utr.bed
**Command 2:** predictAPA -i A1.bedgraph, A2.bedgraph, A3.bedgraph B1.bedgraph, B2.bedgraph, B3.bedgraph -g 2 -n 3 3 -u mm10.utr.bed -o APA_output.txt
**Command 3:** deAPA(‘APA_output.txt’, ‘APA_output.stat.txt’, 1, 2, 1, 1, 20)

### 2 Parameters to run flux-simulator (30 million reads)

| Parameters | Value | Description |
| --- | --- | --- |
| REF_FILE_NAME | mm10.refGene.gtf | GTF reference annotation |
| GEN_DIR | Genome_mm10 | Genomic sequences directory |
| NB_MOLECULES | 8000000 | Number of RNA molecules |
| TSS_MEAN | 100 |  |
| POLYA_SCALE | 100 | Transcript modification parameters |
| POLYA_SHAPE | 2 |  |
| FRAG_SUBSTRATE | DNA |  |
| FRAG_METHOD | NB | Library Preparation parameters |
| FRAG_NB_LAMBDA | 575 |  |
| FRAG_NB_M | 1 |  |
| RTRANSCRIPTION | YES | Switch on reverse transcription |
| PCR_DISTRIBUTION | none |  |
| GC_MEAN | NaN | Amplification parameters |
| GC_SD | NaN |  |
| PCR_PROBABILITY | 0.1 |  |
| FILTERING | YES | Switches size selection On |
| UNIQUE_IDS | TRUE | Create Unique Read Identifiers for paired-end |
| READ_NUMBER | 30000000 | Number of reads |
| READ_LENGTH | 76 | Length of each read |
| PAIRED_END | YES | Paired end reads |
| FASTA | YES | Generate Fasta file |
| ERR_FILE | 76 | Error model for length 76 |

### 3 Realtime quantitative PCR (RT-qPCR) analysis and primer sequences

Total RNAs from TSC1 WT or TSC1-/- MEF cells were isolated by Trizol method according to manufacturer’s protocol (https://assets.thermofisher.com/TFS-Assets/LSG/manuals/trizol_reagent.pdf). Reverse transcription reaction using Oligo-d(T) priming and NxGen M-MuLV Reverse transcriptase (Lucigen) was carried out according to the manufacturer’s protocol (https://www.lucigen.com/docs/manuals/MA115-M-MuLV.pdf). SYBR Green was used to detect and quantitate the PCR products in real-time reactions. Quantitation of the real-time PCR results was done using standard curve method for accuracy and reliability of the analysis. The primer sequences used to measure the RSI for each transcript are as follows:

mRpl22 Total forward 5’-AAGTTCAC CCTGGACTGC AC-3’
mRpl22 Total reverse 5’-GTGATCTT GCTCTTGCTG CG-3’
mRPL22 Long Forward 5’-TGGGCATC TGGGCTTTTA GG-3’
mRPL22 Long reverse 5’-GCTTGTTGCA GACTTGCTCA-3’
mSRSF3 Total forward 5’ - GCTGCCGTGTAAGAGTGGAA-3’
mSRSF3 Total reverse 5’- AGGACTCCTCCTGCGGTAAT-3’
mSRSF3 Long forward 5’ - TGCAACAGTCTTGTGGCTTA-3’
mSRSF3 Long reverse 5’-TGCAATGGCTCTTACATAGACC-3’

### 4 Venn diagram for BT549 mock vs BT549 Torin1 treated

**Figure S1:**
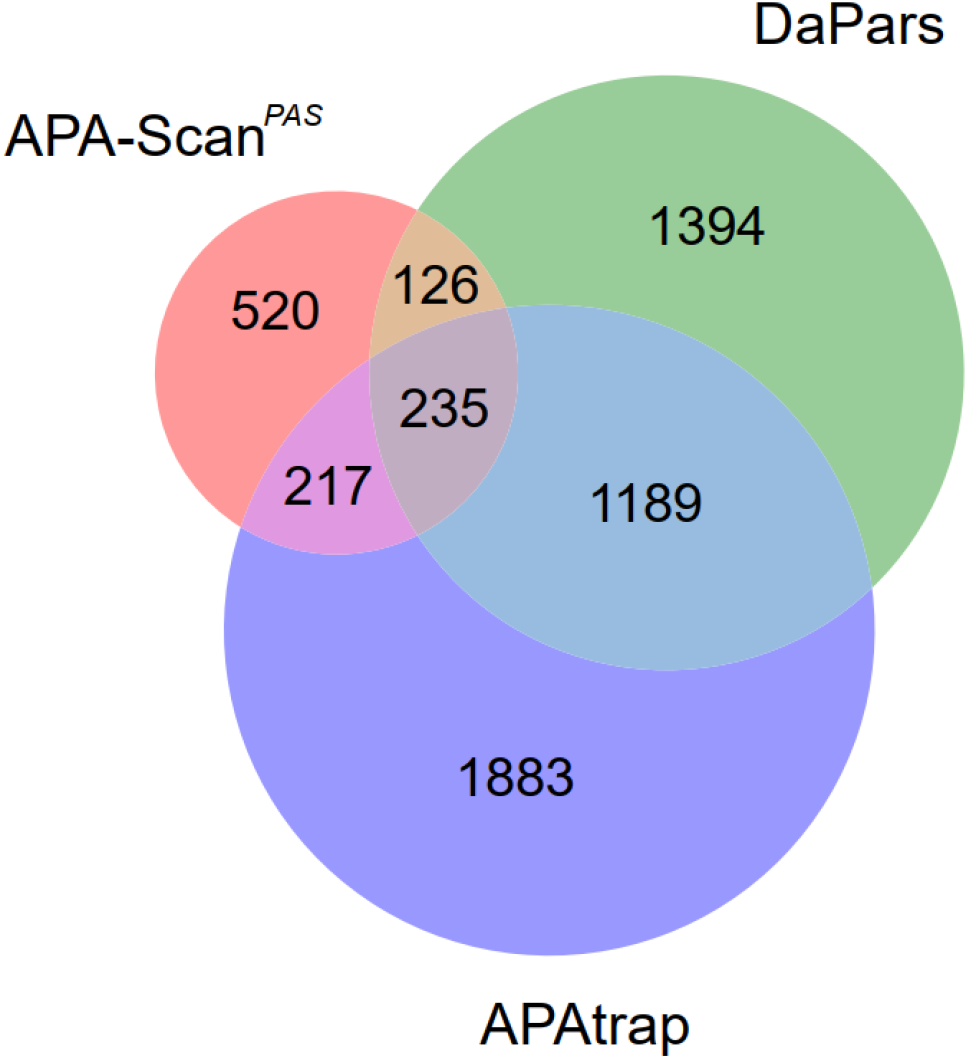
Venn diagram shows the overlapped genes with the 3’-UTR APA events identified by three methods between two breast cancer cell lines (BT549 mock vs BT549 Torin1 treated).

### 5 Experimental results with TCGA BRCA samples

One pair of TCGA breast cancer normal and tumor samples were selected and ran through APA-Scan for the detection of 3’-UTR APA. A total of 1266 APA events were detected between the two samples, whereas 1170 (92%) significant APA events were identified in tumor and 96 (8%) significant APA events were found in normal tissue sample. To inspect the correlation between 3’-UTR APA events and the gene expression profiles, we did the differential gene expression analysis and identified the genes in both tumor and normal samples. The result is illustrated in Figure 1. In the scatter plot, the y-axis denotes the Log_2_ fold-change in the differential gene expression analysis and the x-axis shows the significance of UTR-APA (Log_10_ *p*-value). The left three sections and the right three sections show the 3’-UTR truncated genes in tumor and normal sample, respectively. The top three sections and the bottom three sections represent the up-regulated and down-regulated genes in normal tissue over tumor sample. This plot leads us to the observation that majority (>72%) of the 3’-UTR APA genes are not differentially expressed.

**Figure S2:**
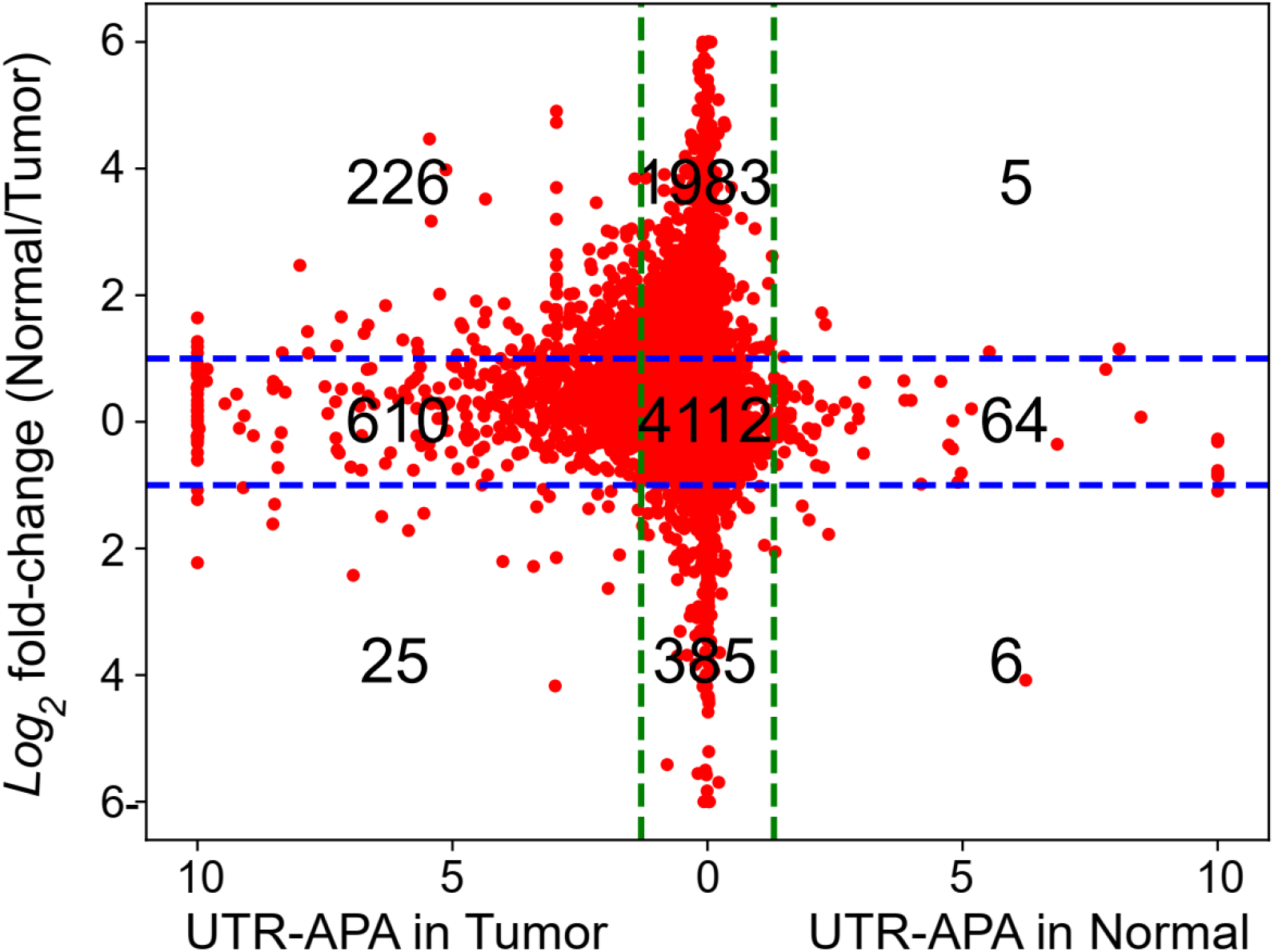
Scatter plot of APA and differentially expressed genes for TCGA tumor (TCGA-BH-A0BQ-01A) vs. matched normal tissue (TCGA-BH-A0BQ-11A) sample. Red dots represent individual gene in the analysis. Horizontal blue-dashed lines represent the cutoff values for two-fold changes in differential gene expression. Vertical green-dashed lines represent the cutoff values for log_10_(p-value) of 3’-UTR APA determined by the Chi-squared test.

## APA-Scan User Manual

### 1 About

APA-Scan is a computational tool which can detect and visualize genome-wide 3’-UTR APA events. APA-Scan integrates both 3’-end-seq (an RNA-seq method with a specific enrichment of 3’-ends ofmRNA) data and the location information of predicted canonical PASs with RNA-seq data to improve the quantitative definition of genome-wide UTR-APA events. It is also advantageous in producing high quality plots of the user defined events.

### 2 Download

APA-Scan is downloadable directly from https://github.com/compbiolabucf/APA-Scan. Users need to have python (version 3.0 or higher) installed in their machine to run APA-Scan.

### 3 Required Softwares

1. Python (version 3.0 or higher)
2. Samtools 0.1.8* [This specific version]

#### Required python packages

1. Pandas: $ pip install pandas
2. Bio: $ pip install biopython
3. Scipy: $ pip install scipy
4. Numpy: $ pip install numpy
5. Peakutils: $ pip install PeakUtils

### 4 Run APA-Scan

APA-Scan can handle both human and mouse data for detecting potential APA truncation sites. The tool is designed to follow the format of Refseq annotation and genome file from UCSC Genome Browser. Users need to have the following two files in the parent directory in order to run APA-Scan:

1. Refseq annotation (.txt format)
2. Genome fasta file (downloaded from UCSC genome browser)

#### 4.1 Required files

APA-Scan has two python scripts: **APA-Scan.py, Make-Plots.py**

And 1 configuration file: **configuration.ini**

The configuration file allows the user to specify the directories of the input samples, the species to be analyzed and the directory where all output files will be stored.

APA-Scan supports the analysis of multiple samples that belong to two different groups-all BAM files inside the input1 directory will be considered as part of the first group, and all BAM files inside the input2 directory will be considered as part of the second group. It is required to have at least one BAM file in each input directory.

#### 4.2 Running with parameters in the configuration.ini file

|  |  |
| --- | --- |
| species*: | Species name (human/mouse) |
| input1* : | Directory containing the first group of samples with RNA-seq data [must be a folder name without '/' at the end] |
| input2* : | Directory containing the second group of samples with RNA-seq data[must be a folder name without '/' at the end] |
| pas1* : | Directory containing the first group of samples with 3'-end-seq data [must be a folder name without '/' at the end]. Default is NULL |
| pas2* : | Directory containing the second group of samples with 3'-end-seq data [must be a folder name without '/' at the end]. Default is NULL |
| extended* : | APA-Scan will run on 'Extended 3UTR' mode and it will search for APA sites upto 10kb downstream of the annotated transcript. Value: yes or no |
| All* : | If selected 'yes', APA-Scan will report all the candidate cleavage sites of a gene, whether they are significant or not. Otherwise, APA-Scan will report the most significant event for each gene [default]. Value: yes or no |
| annotation* | RefSeq annotation file, downloaded from UCSC Genome Browser, in .txt format |
| genome* | Genome fasta file, in .fa format |
| output_dir : | Output directory for writing the results. [optional] |

An example of the congiration.ini file is provided below:

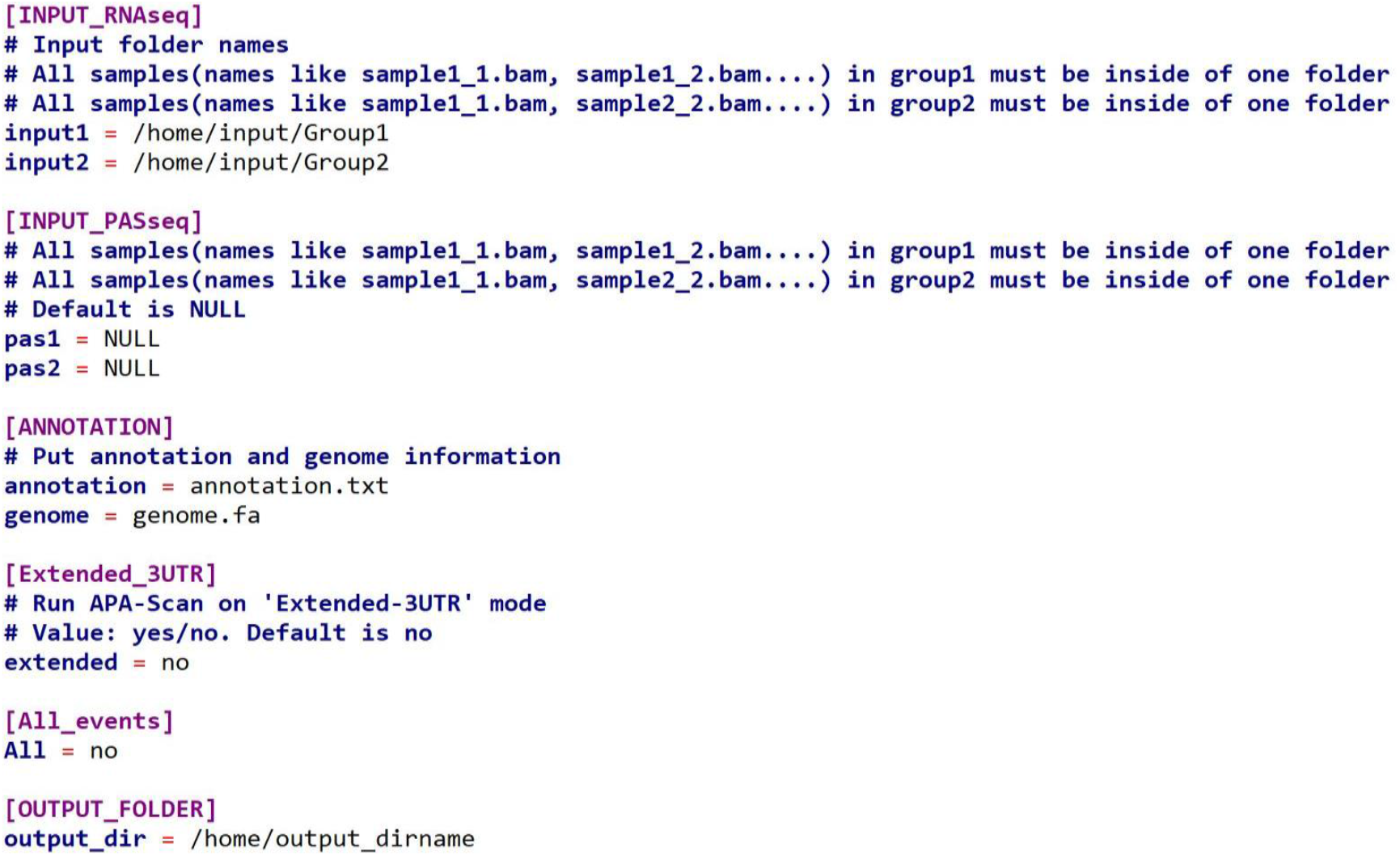

Once the parameters have been specified in the configuration file, the user will open a terminal and enter the following command to run APA-Scan:

##### $ python3 APA-Scan.py

APA-Scan.py will generate several intermediary files in the output directory. After computing the significance of the association between the two groups of samples, the final results will be written in the file named **Group1_Vs_Group2.csv**. The following image shows some of the generated fields in Group1_Vs_Group2.csv:

| Chrom | Gene Name | Strand | Start | End | Position | p-value | Ratio Difference | Absolute ratio difference |
| --- | --- | --- | --- | --- | --- | --- | --- | --- |
| chr19 | RPL13A | + | 49491728 | 49492307 | 49491826 | 1.92E-42 | -0.065610766 | 0.065610766 |
| chr10 | VIM | + | 17237229 | 17237597 | 17237318 | 4.61E-31 | 0.028446051 | 0.028446051 |
| chr17 | RPL26 | - | 8377515 | 8377692 | 8377564 | 2.98E-26 | -0.04344075 | 0.04344075 |
| chr5 | STC2 | - | 173314722 | 173318249 | 173317376 | 2.33E-22 | 0.112542951 | 0.112542951 |
| chr17 | RPL19 | + | 39204524 | 39204730 | 39204650 | 1.03E-21 | -0.049312569 | 0.049312569 |
| chr13 | CDC16 | + | 114272183 | 114272726 | 114272209 | 4.05E-17 | -0.195062219 | 0.195062219 |
| chr7 | HNRNPA2B1 | - | 26189935 | 26192577 | 26191861 | 5.11E-16 | -0.061955041 | 0.061955041 |
| chr9 | RABL6 | + | 136833719 | 136834476 | 136834327 | 5.68E-15 | -0.30708408 | 0.30708408 |
| chrX | RPS4X | - | 72272602 | 72272772 | 72272628 | 2.08E-14 | 0.041582181 | 0.041582181 |
| chr1 | STMN1 | - | 25900115 | 25901087 | 25900534 | 2.9E-14 | 0.043951567 | 0.043951567 |
| chr17 | RPAIN | + | 5432541 | 5433020 | 5432863 | 3.79E-14 | 0.061525048 | 0.061525048 |
| chr6 | SOD2 | - | 159679063 | 159682638 | 159682113 | 8.77E-14 | -0.076980538 | 0.076980538 |
| chr11 | CCDC84 | + | 119015537 | 119015792 | 119015726 | 2.19E-12 | -0.057536734 | 0.057536734 |
| chr1 | SYNC | - | 32679905 | 32681860 | 32680315 | 4.57E-12 | 0.095614951 | 0.095614951 |
| chr1 | RPS8 | + | 44778575 | 44778740 | 44778687 | 1.21E-11 | 0.037494068 | 0.037494068 |
| chr8 | SFRP1 | - | 41261956 | 41265489 | 41265472 | 5.35E-11 | -0.292512141 | 0.292512141 |
| chr15 | TPM1 | + | 63069869 | 63071914 | 63069945 | 5.54E-11 | 0.075237838 | 0.075237838 |
| chr1 | MEF2D | - | 156463720 | 156467656 | 156467650 | 1.5E-10 | -0.282933192 | 0.282933192 |
| chr5 | PHYKPL | - | 178208473 | 178211970 | 178211572 | 3.28E-10 | 0.079757735 | 0.079757735 |
| chr2 | RPL37A | + | 216501340 | 216501465 | 216501446 | 3.91E-10 | 0.019741659 | 0.019741659 |
| chr12 | PRIM1 | - | 56731579 | 56731734 | 56731638 | 8.48E-10 | 0.190879674 | 0.190879674 |
| chr18 | RMC1 | + | 23531624 | 23531807 | 23531765 | 9.45E-10 | -0.13019906 | 0.13019906 |

### 5 Run Make-plots.py

Make-plots.py also requires the same configuration file to run. It will use the input and output directories listed in the configuration file and prepare a read coverage plot along with the 3’-UTR annotation based on user defined region.

#### python3 Make-plots.py

After executing this command above for a few seconds, Make-plots.py will ask the user to insert the region of interest in a specific format:

#### Chrom:GeneName:RegionStart-RegionEnd

#### 5.1 Make-plots.py parameter descriptions

|  |  |
| --- | --- |
| Chrom | Name of the chromosome |
| GeneName | Name of the gene |
| RegionStart | Starting position of the region |
| Region End | End position of the region |

Example: **chr1:Tceb1:16641724-16643478**

Make-Plots.py will generate a visual representation of the results shown for each of the regions entered. The plot will illustrate the most significant transcript cleavage site with a red vertical bar on top of RNA-seq read data (and 3’end-seq if available). If the input parameters have 3’end-seq information along with the RNA-seq, then it will generate plots for both cases (See figure below). It will also show the UTR truncation point (annotated and unannotated) at the bottom panel.

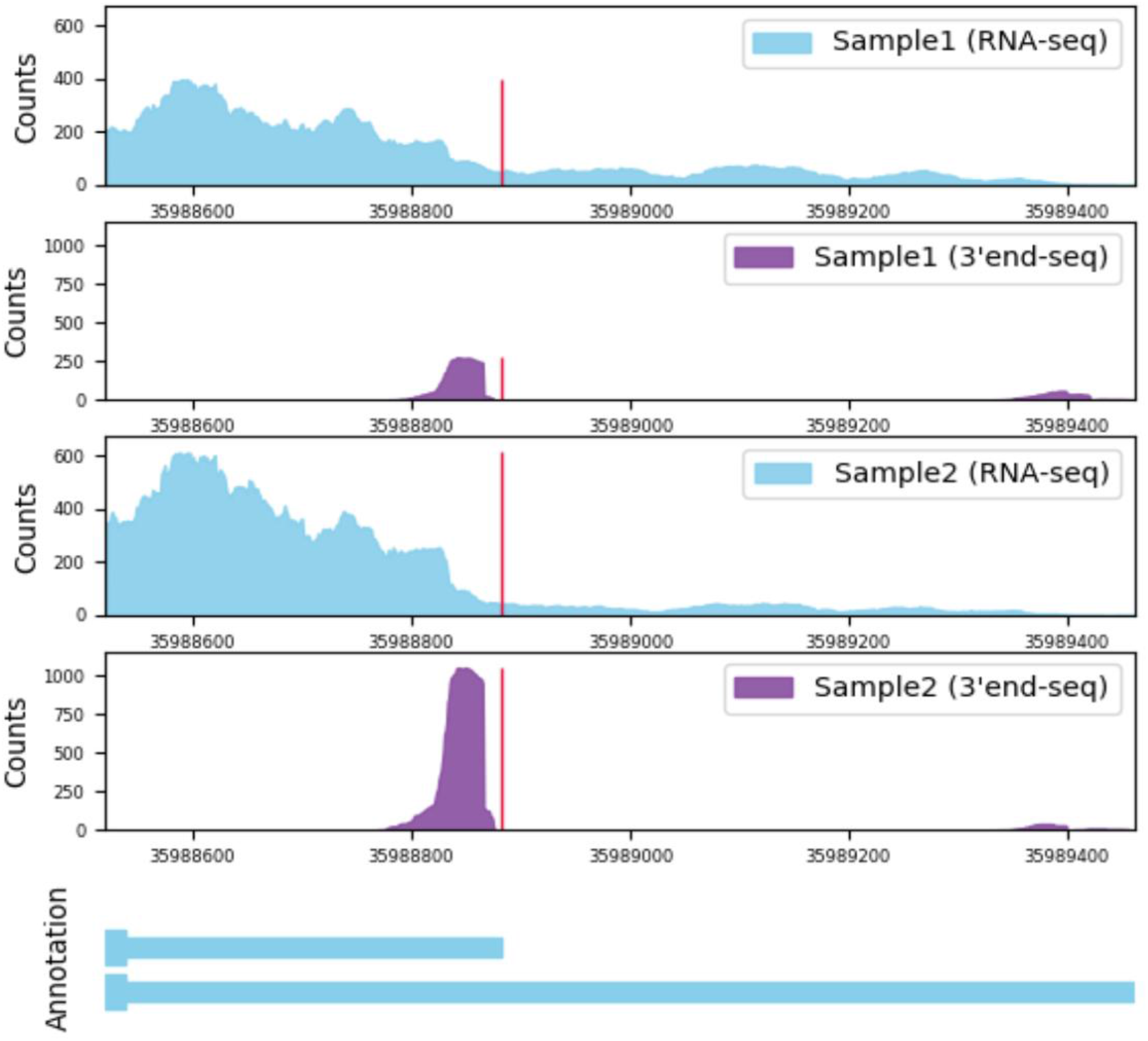

The first two subplots of the figure represent the read coverage of the two biological conditions. The bottom subplot shows the gene annotation and the exon information of that gene.

## References

1. Proudfoot, N.J.: Ending the message: poly (A) signals then and now. Genes & development 25(17), 1770–1782 (2011)

2. Tian, B., Manley, J.L.: Alternative cleavage and polyadenylation: the long and short of it. Trends in Biochemical Sciences 38(6), 312–320 (2013)

3. Elkon, R., Ugalde, A.P., Agami, R.: Alternative cleavage and polyadenylation: extent, regulation and function. Nature Reviews Genetics 14(7), 496 (2013)

4. Yeh, H.-S., Zhang, W., Yong, J.: Analyses of alternative polyadenylation: from old school biochemistry to high-throughput technologies. BMB reports 50(4), 201 (2017)

5. Mayr, C., Bartel, D.P.: Widespread shortening of 3’ UTRs by alternative cleavage and polyadenylation activates oncogenes in cancer cells. Cell 138(4), 673–684 (2009)

6. Lembo, A., Di Cunto, F., Provero, P.: Shortening of 3 UTRs correlates with poor prognosis in breast and lung cancer. PloS one 7(2), 31129 (2012)

7. Morris, A.R., Bos, A., Diosdado, B., Rooijers, K., Elkon, R., Bolijn, A.S., Carvalho, B., Meijer, G.A., Agami, R.: Alternative cleavage and polyadenylation during colorectal cancer development. Clinical Cancer Research 18(19), 5256–5266 (2012)

8. Chang, J.-W., Zhang, W., Yeh, H.-S., De Jong, E.P., Jun, S., Kim, K.-H., Bae, S.S., Beckman, K., Hwang, T.H., Kim, K.-S., et al.: mRNA 3-UTR shortening is a molecular signature of mTORC1 activation. Nature communications 6(1), 1–9 (2015)

9. Chang, J.-W., Zhang, W., Yeh, H.-S., Park, M., Yao, C., Shi, Y., Kuang, R., Yong, J.: An integrative model for alternative polyadenylation, IntMAP, delineates mTOR-modulated endoplasmic reticulum stress response. Nucleic Acids Research 46(12), 5996–6008 (2018)

10. Hoffman, Y., Bublik, D.R., P. Ugalde, A., Elkon, R., Biniashvili, T., Agami, R., Oren, M., Pilpel, Y.: 3’UTR shortening potentiates microRNA-based repression of pro-differentiation genes in proliferating human cells. PLoS genetics 12(2), 1005879 (2016)

11. Sandberg, R., Neilson, J.R., Sarma, A., Sharp, P.A., Burge, C.B.: Proliferating cells express mRNAs with shortened 3’untranslated regions and fewer microRNA target sites. Science 320(5883), 1643–1647 (2008)

12. Xia, Z., Donehower, L.A., Cooper, T.A., Neilson, J.R., Wheeler, D.A., Wagner, E.J., Li, W.: Dynamic Analyses of Alternative Polyadenylation from RNA-Seq Reveal 3’-UTR Landscape Across 7 Tumor Types. Nature Communications 5, 5274 (2014)

13. Wang, W., Wei, Z., Li, H.: A change-point model for identifying 3’ UTR switching by nextgeneration RNA sequencing. Bioinformatics 30(15), 2162–2170 (2014)

14. Le Pera, L., Mazzapioda, M., Tramontano, A.: 3USS: a web server for detecting alternative 3’ UTRs from RNA-seq experiments. Bioinformatics 31(11), 1845–1847 (2015)

15. Ye, C., Long, Y., Ji, G., Li, Q.Q., Wu, X.: APAtrap: identification and quantification of alternative polyadenylation sites from RNA-seq data. Bioinformatics 34(11), 1841–1849 (2018)

16. Magana-Mora, A., Kalkatawi, M., Bajic, V.B.: Omni-PolyA: a method and tool for accurate recognition of Poly(A) signals in human genomic DNA. BMC genomics 18(1), 1–13 (2017)

17. Shepard, P.J., Choi, E.-A., Lu, J., Flanagan, L.A., Hertel, K.J., Shi, Y.: Complex and dynamic landscape of rna polyadenylation revealed by pas-seq. Rna 17(4), 761–772 (2011)

18. Griebel, T., Zacher, B., Ribeca, P., Raineri, E., Lacroix, V., Guigó, R., Sammeth, M.: Modelling and simulating generic RNA-Seq experiments with the flux simulator. Nucleic acids research 40(20), 10073–10083 (2012)

19. Oshlack, A., Robinson, M.D., Young, M.D.: From RNA-seq reads to differential expression results. Genome biology 11(12), 220 (2010)

20. Gruber, A.J., Schmidt, R., Gruber, A.R., Martin, G., Ghosh, S., Belmadani, M., Keller, W., Zavolan, M.: A comprehensive analysis of 3 end sequencing data sets reveals novel polyadenylation signals and the repressive role of heterogeneous ribonucleoprotein C on cleavage and polyadenylation. Genome research 26(8), 1145–1159 (2016)

21. Li, H., Handsaker, B., Wysoker, A., Fennell, T., Ruan, J., Homer, N., Marth, G., Abecasis, G., Durbin, R.: The sequence alignment/map format and SAMtools. Bioinformatics 25(16), 2078–2079 (2009)

22. Kim, D., Pertea, G., Trapnell, C., Pimentel, H., Kelley, R., Salzberg, S.L.: TopHat2: accurate alignment of transcriptomes in the presence of insertions, deletions and gene fusions. Genome biology 14(4), 1–13 (2013)

23. Langmead, B., Trapnell, C., Pop, M., Salzberg, S.L.: Ultrafast and memory-efficient alignment of short DNA sequences to the human genome. Genome biology 10(3), 1–10 (2009)

